# In the back of your mind: Cortical mapping of paraspinal afferent inputs

**DOI:** 10.1101/2021.12.19.473341

**Authors:** David M. Cole, Philipp Stämpfli, Robert Gandia, Louis Schibli, Sandro Gantner, Philipp Schuetz, Michael L. Meier

**Affiliations:** Department of Chiropractic Medicine, Integrative Spinal Research, Balgrist University Hospital, University of Zurich, Zurich, Switzerland; Department of Psychiatry, Psychotherapy and Psychosomatics, Hospital of Psychiatry, University of Zurich, Zurich, Switzerland; MR-Center of the Psychiatric Hospital, University of Zurich, Zurich, Switzerland; Competence Center Thermal Energy Storage, Lucerne University of Applied Sciences and Arts, Lucerne, Switzerland

**Keywords:** fMRI, somatosensory, cortex, proprioception, back, somatotopy, motor control

## Abstract

Topographic organization is a hallmark of vertebrate cortex architecture, characterized by ordered projections of the body’s sensory surfaces onto brain systems. High-resolution functional magnetic resonance imaging (fMRI) has proven itself as a valuable tool to investigate the cortical landscape and its (mal-)adaptive plasticity with respect to various body part representations, in particular extremities such as the hand and fingers. Less is known, however, about the cortical representation of the human back. We therefore validated a novel, MRI-compatible method of mapping cortical representations of sensory afferents of the back, using vibrotactile stimulation at varying frequencies and paraspinal locations, in conjunction with fMRI. We expected high-frequency stimulation to be associated with differential neuronal activity in the primary somatosensory cortex (S1) compared to low-frequency stimulation and that somatosensory representations would differ across the thoracolumbar axis. We found significant differences between neural representations of high- and low-frequency stimulation and between representations of thoracic and lumbar paraspinal locations, in several bilateral S1 sub-regions, and in regions of the primary motor cortex (M1). High-frequency stimulation preferentially activated Brodmann Area (BA) regions BA3a and BA4p, while low-frequency stimulation was more encoded in BA3b and BA4a. Moreover, we found clear topographic differences in S1 for representations of the upper and lower back during high-frequency stimulation. We present the first neurobiological validation of a method for establishing detailed cortical maps of the human back, which might serve as a novel tool to evaluate the pathological significance of neuroplastic changes in clinical conditions such as chronic low back pain.

**Key points:**

- Detailed investigations of cortical representations of somatosensory paraspinal afferents along the thoracolumbar axis are lacking.
- Using fMRI combined with a novel vibrotactile stimulation device (“pneuVID”) we investigated different sensorimotor cortical representations of the back and explored topographic differences between the upper and lower back.
- We found differential sub-regional sensorimotor neural representations of high- and low-frequency stimulation, as well as revealing initial evidence of the somatotopy of upper and lower paraspinal representations.
- The current approach might serve as a promising tool to elucidate the role of cortical reorganisation in the pathophysiology of clinical conditions such as chronic low back pain.

## 1. Introduction

A principal feature of the organization of the vertebrate brain is its orderly arrangement, characterized by topographic maps that reflect the spatially-preserved relationship between neurons of a source region and neurons of a target region (e.g., neighbouring cells in the skin project to neighbouring cortical neurons; Thivierge and Marcus, 2007). Topographic maps play an important role in the transfer of sensory information and have been studied extensively in human imaging experiments to reveal the somatotopic organization of various body parts, particularly the hand and the fingers (Maldjian et al., 1999; Hluštík et al., 2001; Thivierge and Marcus, 2007; Nelson and Chen, 2008; Schweizer et al., 2008; Ann Stringer et al., 2014; Martuzzi et al., 2014; Kolasinski et al., 2016; Akselrod et al., 2017; Schellekens et al., 2021). However, less is known about the cortical topographic organization of the human back. In his pioneering work, Penfield identified the hip and the shoulder on the convexity of the postcentral gyrus and drew the back between these two areas on the sensory Homunculus (Penfield, 1947). In 2018, intra-cortical stimulation of Brodmann area 1 (BA1) in the primary somatosensory cortex (S1) identified the representations of the thorax and abdomen to indeed lie between the hip and shoulder (Roux et al., 2018). Nonetheless, detailed cortical topographic maps of paraspinal sensory input along the thoracolumbar axis are still lacking, perhaps due to technical challenges in providing reliable sensory stimulation across the back in a magnetic resonance imaging (MRI) environment. Establishing finely-grained cortical maps of the back is not only of interest for fundamental neuroscience but could also be crucial for further research into possible links between brain changes and the development and maintenance of neurological disorders and clinical conditions such as chronic low back pain (LBP; Van Dieën et al., 2017; Schmid et al., 2021).

We here sought to validate a technique combining vibrotactile stimulation of multiple paraspinal locations along the thoracolumbar axis, using a novel pneumatic vibration device (“pneuVID”; Schibli et al., 2021) in conjunction with high-resolution brain functional MRI (fMRI), in order to investigate such key questions of the cortical topography of paraspinal sensory inputs, which hitherto remain unanswered. Although MR-compatible vibration devices exist (Montant et al., 2009; Goossens et al., 2016), this is the first apparatus designed specifically for paraspinal muscle vibration at different segmental levels in the MRI environment. To explore possible distinct contributions of various paraspinal tissue mechanoreceptors to cortical activity, different vibration frequencies were applied. Although a perfectly separated provocation of signal transmission between the different types of mechanoreceptors is not achievable using vibration, lower vibration frequencies (between 5-50 Hz) have been shown to mainly activate Meissner’s corpuscles (located superficially within the dermal papillae), whereas more deeply located mechanoreceptors, such as Pacinian corpuscles or muscle spindles, respond to higher frequencies (between 80-400 Hz, in combination with appropriate stimulation amplitudes – between 0.5-1mm to optimally stimulate muscle spindles; Sato, 1961; Talbot et al., 1968; Weerakkody et al., 2007; Chung et al., 2013; Goossens et al., 2016; Schellekens et al., 2021). Vibrotactile stimulation at different frequencies has, therefore, often been used to decipher the contribution of different mechanoreceptor types to central processing (Harrington and Hunter Downs, 2001; Avanzino et al., 2014; Kim et al., 2016; Schellekens et al., 2021). The central processing of vibrotactile stimuli seems to occur largely in S1, with some evidence alluding to mechanoreceptor-specific and hierarchical processing in the respective Brodmann areas BA3a/3b, BA1 and BA2 (Delhaye et al., 2018; Schellekens et al., 2021). Hence, to evaluate the ability of pneuVID to evoke neural activity characteristic of differential mechanoreceptor contributions and to simultaneously gain initial insights into the topographic organization of the back, we hypothesised that: (i) high-frequency pneuVID stimulation (80 Hz vibration) would activate different somatosensory representations of paraspinal afferent input compared to low-frequency stimulation (20 Hz vibration); and (ii) somatosensory representations along the thoracolumbar axis of the back would be differentiable, providing a promising basis towards discovering a more finely-grained topographic map in S1 than has previously been attainable.

## 2. Methods

### 2.1. pneuVID

#### 2.1.1. pneuVID system

PneuVID was used to apply vibrotactile stimulation to different thoracolumbar segments. PneuVID was specifically designed for human paraspinal tissue vibration in the MRI environment. Vibrotactile stimulation is performed with manually-manufactured vibration units made of silicone and with compressed air pulses operating at a feed pressure of 1.5 bar. The compressed air pulses are controlled by a valve box attached to the end of the MRI scanner bed. The interaction with the MRI system and the operator is provided by a control module (Raspberry Pi, Cambridge, UK) connected via fibre-optic connection to the valve box (Fig. 1). Details about the design, the different system components, and integration of pneuVID in the MR environment are described elsewhere (Schibli et al., 2021). In the current study, two different frequencies (20 Hz, 80 Hz) with a constant feed pressure of 1.5 bar were applied with amplitudes between 0.5-1 mm at each segmental level (see *section 2.2.2*; see also Schibli et al., 2021 for a detailed evaluation of practical considerations regarding amplitude variability).

**Figure 1.**
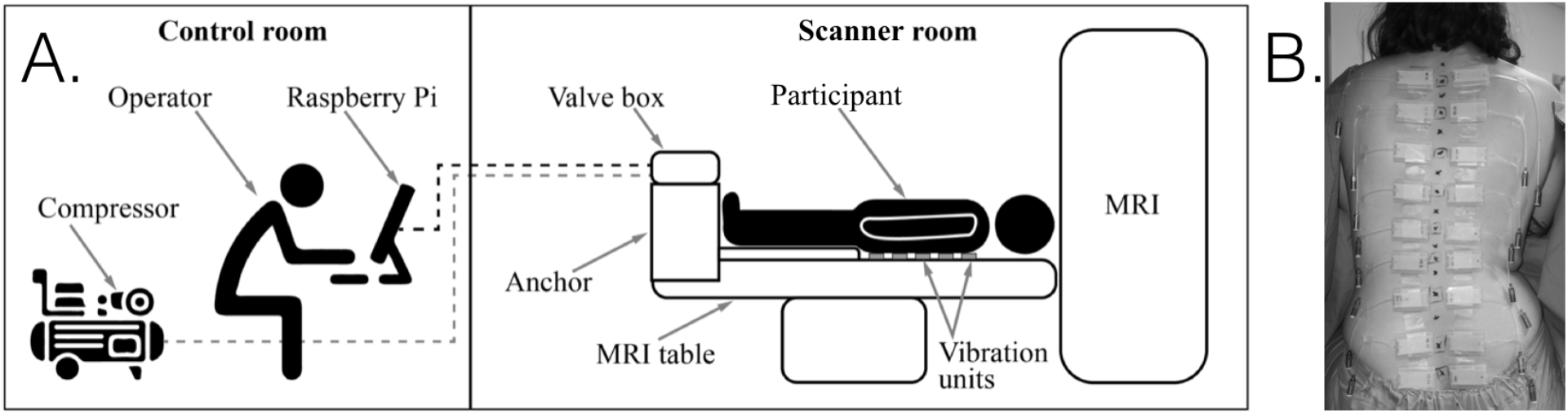
(A) Simplified representation of the pneuVID hardware setup (see also Schibli et al., 2021). Via this system, the operator can safely control the vibrotactile stimulation experiment from outside the MRI scanner room. (B) Example image showing placement of pneuVID vibration units on the skin of the back (corresponding, from top to bottom, to the thoracic, lumbar and sacral spinous processes T3, T5, T7, T9, T11, L1, L3, L5 and sacral[S1]).

### 2.2. Neuroimaging methods

#### 2.2.1 Participants

Fifteen healthy volunteers (7 female; mean age 29.9 years, SD ± 5.1) were recruited and underwent fMRI scanning. Exclusion criteria were excessive consumption of alcohol or consumption of other drugs or analgesics within the last 24 h, prior foot or ankle surgery, any neuromuscular diseases that might affect gait and posture, injuries of the motor system with permanent deformities and body mass index (BMI) > 30 kg/m^2^. The study was approved by the Ethics Committee Zurich (Switzerland, ID 2018-01001) and all participants provided written informed consent before participation. The study was conducted in accordance with the Declaration of Helsinki.

#### 2.2.2. Paradigm

Prior to the attachment of the pneuVID units, nine different spinous processes (T3, T5, T7, T9, T11, L1, L3, L5 and sacral[S1]) were marked through palpation by an experienced physiotherapist. Twenty pneuVID vibrotactile stimulation units were applied bilaterally to the nine paraspinal locations along the erector spinae muscles (Fig. 1B) and one on the calves of the legs at the triceps surae muscles (mTS), using adhesive tape and a sugar-based glue that was developed in-house. These stimulators were activated, always bilaterally at a single one of the ten locations, at either 20 Hz or 80 Hz vibration frequency, in a pseudorandomised order (no more than three occasions in a row at the same location or frequency) that differed per run and per participant. Stimulation events were always of five seconds duration and the inter-stimulation interval was jittered within the range of 4-7 s. A total of 120 stimulation events were delivered per run (12 per location, further divided into 6 per frequency type per location). Two separate runs of the pneuVID stimulation paradigm were delivered to each participant during fMRI data acquisition. Prior to the experimental runs, all participants underwent a descending test vibration sequence where they consistently reported feeling stimulation at each location in the midline of the back (i.e., they perceived no lateralisation of stimulation) and at the mTS and could distinguish perceptions evoked by 80 Hz and 20 Hz at each of these locations.

#### 2.2.3. Image acquisition

Neuroimaging data were acquired using a 3 T Philips Achieva MRI scanner with a 32-channel sensitivity encoding (SENSE)-compatible head coil (Philips, Best, The Netherlands) at the MRI Center of the Psychiatric Hospital, University of Zurich (PUK). The stimulation paradigm was separated into two runs of 22.5 minutes length each. We acquired 750 volumes of T2*-weighted echo-planar imaging (EPI) data sensitive to blood-oxygenation level-dependent (BOLD) contrast during both paradigm runs (TR = 1.8 s; TE = 34 ms; flip angle = 70°; 54 interleaved ascending axial slices with multiband factor = 3; in-plane resolution = 1.72 × 1.72 mm; slice thickness = 2.0 mm with 0 mm slice gap; SENSE factor = 1.4; EPI factor = 87). The scanner automatically acquired and removed initial dummy volumes for magnetic equilibration. Before the second stimulation run, we also acquired a high-resolution anatomical T1-weighted (T1w) volume (MPRAGE; TR = 6.6 ms; TE = 3.1 ms; flip angle = 9°; field-of view 230 × 226 × 274; voxel size = 1.0 × 1.0 × 1.2 mm; turbo field echo factor = 203).

#### 2.2.4. Image preprocessing

Prior to preprocessing, the neuroimaging data were converted to the consensus Brain Imaging Data Structure (BIDS) format (Gorgolewski et al., 2016) and then subjected to preliminary assessment using the *FMRIprep* pipeline (v. 20.2.0) (Esteban et al., 2018). As part of this procedure, each T1w volume was corrected for intensity non-uniformity and skull-stripped (using the OASIS template) using Advanced Normalization Tools (ANTs) v2.1.0 (Tustison et al., 2010). Brain surfaces were reconstructed using *recon-all* from FreeSurfer v6.0.1 (Dale et al., 1999), and the brain mask estimated previously was refined with a custom variation of the method to reconcile ANTs-derived and FreeSurfer-derived segmentations of the cortical grey-matter of *Mindboggle* (Klein et al., 2017). Spatial normalization to the ICBM 152 Nonlinear Asymmetrical (MNI) template 6^th^ generation version was performed through nonlinear registration with ANTs v2.1.0 (Avants et al., 2008), using brain-extracted versions of both T1w volume and template. Brain tissue segmentation of cerebrospinal fluid (CSF), white-matter (WM) and grey-matter (GM) was performed on the brain-extracted T1w using FSL (v5.0.9) FAST (Zhang et al., 2001).

Functional data were motion corrected using the MCFLIRT tool of FSL (Smith et al., 2004). This was followed by co-registration with six degrees of freedom to the corresponding T1w image using boundary-based registration (Greve and Fischl, 2009). Motion correcting transformations, BOLD-to-T1w transformation and T1w-to-template (MNI) warp were concatenated and applied in a single step using ANTs v2.1.0 (using Lanczos interpolation).

We selected mean time series, their squared versions and the first derivatives of both of these from CSF and WM tissue sources separately, as calculated by *FMRIprep*, for use as confounding signals with which to filter the fMRI data (NaN values at the beginning of the derivative vectors were instead computed as the mean value of the signal vector, in order to allow for regression). In addition to filtering for these eight confound vectors, we also regressed out the time series from all components in the data identified as (primarily motion) artefacts by ICA-based Automatic Removal Of Motion Artifacts (ICA-AROMA; Pruim et al., 2015). This regression filtering of tissue-based and ICA-based identified artefactual noise was performed in a single step, without any temporal filtering (of the data or the confound regressors), on the unsmoothed, MNI-space-normalized output of *FMRIprep*, prior to any participant-level statistical analyses, using *fsl_regfilt*. The outputs from this step thus comprised “aggressively denoised” (Pruim et al., 2015) fMRI datasets from the stimulation paradigm runs.

#### 2.2.5. Statistical analysis

The denoised datasets from the previous processing stage were analysed per run at the first-/participant-level using FSL FEAT (Version 6.00) with FILM prewhitening enabled (Smith et al., 2004). For this initial analysis step, a high pass temporal filter (100 s) was applied to both the fMRI data and the simulation paradigm model design, while the data were also smoothed using a 4 mm full width at half maximum (FWHM) Gaussian kernel. Base regressors for each of the 20 stimulation settings (10 locations × two frequency settings) were modelled as separate explanatory variables (EVs). These 20 EVs were defined per run in terms of the precise onsets of each of the six stimulation events per frequency and location for a given setting, with their durations (5 s) fully modelled, using a haemodynamic response function with Gamma convolution. General linear model (GLM) contrasts of interest were computed including comparisons of: (i) 80 Hz > 20 Hz across all nine paraspinal locations (the mTS leg units were omitted); (ii) the inverse 20 > 80 Hz contrast; (iii) frequency-specific contrasts of the upper four pneuVID units (T3, T5, T7, T9) > lower four panels (L1, L3, L5 and sacral[S1]); and (iv) the inverse 4 lower > 4 upper contrast. In addition, we examined contrasts for individual EV (location per frequency) representations, in order to explore potentially more fine-grained neural representations, in particular of high-frequency sensory stimulatory input. In an attempt to further improve the distinct spatial visualisation of these latter statistical representations, we used FreeSurfer tools to additionally portray versions of these maps on a cortical surface projection corresponding to an ‘inflated’ version of MNI standard space.

Outputs from first-level analyses were subjected to a second, mid-level analysis at the level of individual participants, wherein contrast parameter estimates from the two runs of each participant were tested for their mean representation at the level of fixed effects (using FSL FEAT).

Finally, the resulting averaged participant-level contrasts of interest were entered into mixed effects-level one-sample *t*-tests at the group level using FSL FEAT. The resulting statistical maps of group-level parametric effects for the four contrasts of interest described above were thresholded for significance using whole-brain, cluster-level (*Z* > 2.3; p < 0.05) family-wise error (FWE) correction for testing across multiple voxels (for additional statistical context and interpretation, we also present these results alongside those established using a more conservative cluster-level FWE-corrected threshold: *Z* > 2.81; p < 0.05).

## 3. Results

We found BOLD fMRI activation to be significantly increased under 80 Hz, relative to 20 Hz vibrotactile stimulation of the back in several brain regions (Fig. 2; Table 1). These significant clusters were located in bilateral primary somatosensory and motor cortical regions, as well as right supramarginal gyrus extending into posterior insula, secondary somatosensory cortex and temporoparietal junction regions. More fine-grained spatial analysis of these three activation clusters located in motor and somatosensory cortical regions that were central to our hypotheses, revealed the peak voxel activations to be located, based on their correspondence with the Juelich Histological Atlas (Geyer et al., 1996, 2000): with highest probability for Cluster “Index 1” (Table 1) in the left primary somatosensory cortex S1 (region BA3a 29 %, with 10 % probability of location in BA3b) and 15 % in primary motor cortex M1 (BA4p); for Cluster Index 2 in right BA3a (67 %, with 10 % probability of location in BA3b) and 18% in BA4p; and for Cluster Index 3 in right posterior supramarginal gyrus (11 %). More detailed cluster information is provided in Table 1.

**Figure 2.**
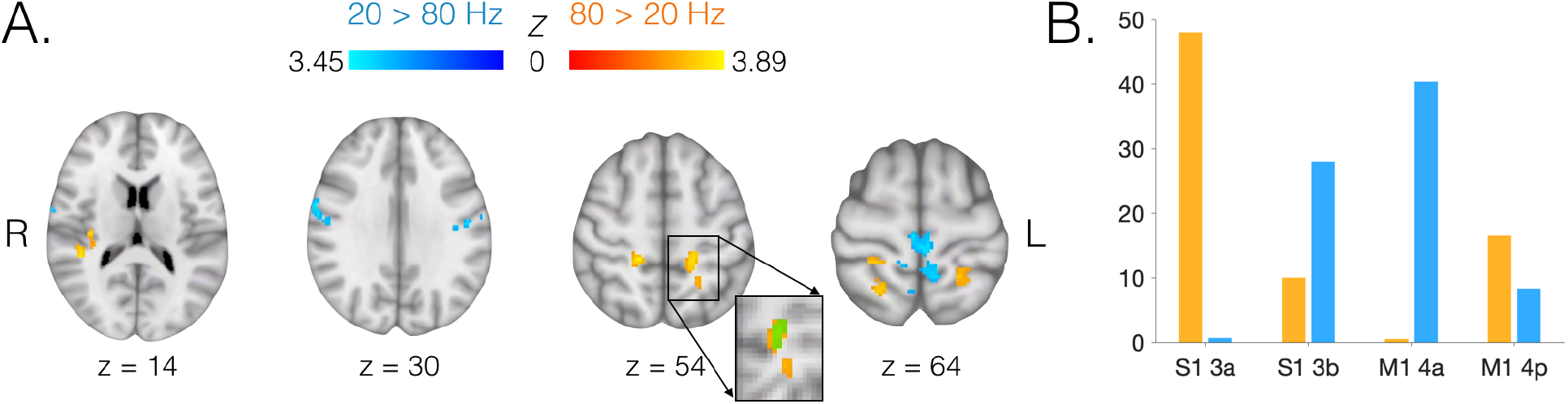
(A) Brain regions displaying differential activation under high- and low-frequency paraspinal vibrotactile stimulation (80 Hz vs. 20 Hz). Red-yellow *Z*-statistic map and colour bar denote 80 Hz > 20 Hz high-frequency-specific significant activation effects (cluster *Z* > 2.3; p < 0.05; FWE-corrected; inset: green overlay with reduced opacity denotes significant equivalent cluster at a more conservative threshold of *Z* > 2.81); blue-cyan denotes low-frequency (20 Hz > 80 Hz) stimulation effects. Statistical maps are overlaid on a background image of a T1 template in MNI152 standard space. R = right hemisphere; L = left. Axial slices are displayed in line with radiological orientation conventions (L/R hemisphere flipped in figure). NB: Nearby regions of significant activation for a given contrast can appear spatially distinct on the representative slices chosen for visualisation but still form part of the same spatially contiguous cluster. (B) Mean % probability (y-axis) across relevant clusters from (A) spanning regions S1 BA3 and/or M1 BA4, in terms of the Juelich Histological Atlas-based likelihood of cluster peaks being located in sub-regions S1 BA3a, S1 BA3b, M1 BA4a and M1 BA4p; a preferential effect can be seen for proprioception-related stimulation (80 > 20 Hz, orange) representation in BA3a, relative to BA3b and in BA4p, relative to 4a, with an apparent opposing preferential effect for non-proprioceptive tactile stimulation (20 > 80 Hz, blue) representation in M1 BA4a, relative to BA4p and S1 BA3b, relative to BA3a.

**Table 1.** Brain regions significantly more activated under 80 Hz, relative to 20 Hz, vibrotactile stimulation of the back. Three regional statistical maps displayed significantly greater activation (cluster *Z* > 2.3; p < 0.05; FWE-corrected; values in parentheses denote significant equivalent cluster information at a more conservative threshold of *Z* > 2.81) during 80 Hz stimulation than during 20 Hz simulation. Statistical and anatomical co-ordinate information (relative to the MNI152 standard space template) is provided. R = right hemisphere; L = left hemisphere; M1 = primary motor cortex; S1 = primary somatosensory cortex; S2 = secondary somatosensory cortex; SPL = superior parietal lobule; TPJ = temporoparietal junction; BA = Brodmann area.

| Cluster Index | Voxels | P | Z-max | x (mm) | y (mm) | z (mm) | Anatomical loci |
| --- | --- | --- | --- | --- | --- | --- | --- |
| 1 | 241<br>(70) | < 0.001<br>(0.006) | 3.89 | -16 | -36 | 52 | L S1 BA3a & BA3b/M1<br>BA4p/SPL |
| 2 | 191 | < 0.001 | 3.85 | 16 | -36 | 54 | R S1 BA3a & BA3b/M1<br>BA4p/SPL |
| 3 | 111 | 0.040 | 3.68 | 44 | -38 | 14 | R supramarginal<br>gyrus/S2/TPJ/posterior insula |

In addition to greater activation observed under high-frequency stimulation, we found some brain regions to be significantly more activated by the lower frequency 20 Hz, relative to 80 Hz, stimulation (Fig. 2A; Table 2). These cluster peak voxels were located, firstly, at the midline in bilateral primary motor and somatosensory cortex, specifically in M1 BA4a (85 %, with only 20 % probability of location in M1 BA4p) and, secondly and thirdly, bilaterally in primary somatosensory cortex, predominantly in BA3b (left 43 %, right 41%). More detailed cluster information is provided in Table 2.

**Table 2.** Brain regions significantly more activated under 20 Hz, relative to 80 Hz, vibrotactile stimulation of the back. Three regional statistical maps displayed significantly greater activation (cluster *Z* > 2.3; p < 0.05) during 20 Hz stimulation than during 80 Hz simulation. Statistical and anatomical co-ordinate information (relative to the MNI152 standard space template) is provided. R = right hemisphere; L = left hemisphere; M1 = primary motor cortex; S1 = primary somatosensory cortex; SPL = superior parietal lobule; BA = Brodmann area.

| Cluster Index | Voxels | P | Z-max | x (mm) | y (mm) | z (mm) | Anatomical loci |
| --- | --- | --- | --- | --- | --- | --- | --- |
| 1 | 293 | < 0.001 | 3.45 | -6 | -30 | 70 | Bilateral M1 BA4a/S1<br>BA3b/SPL |
| 2 | 127 | 0.017 | 3.32 | 62 | 0 | 28 | R S1 BA3b &<br>BA1/BA6/M1 BA4a |
| 3 | 114 | 0.045 | 3.52 | -58 | -12 | 36 | L S1 BA3b & BA2/BA1/M1<br>BA4a |

Overall, these described frequency-dependent results appeared to pursue a pattern of differential representations in key sub-regions of both S1 and M1, seemingly in an opposing manner for the two frequency contrasts. Therefore, we further amalgamated the atlas-based probabilistic information of the relevant cluster activation peak voxels (i.e., Table 1 Cluster Indices 1 and 2; Table 2 Cluster Indices 1-3) in order to permit a simple graphical representation of how the sub-regional dominances of BA3a *vs* BA3b and BA4a *vs* BA4p differed across stimulation frequencies (Fig. 2B).

Finally, we tested for differences between cortical representations of the thoracic (upper) and lumbar including sacral[S1] (lower) paraspinal stimulation, while also exploring more fine-grained representational variability along the thoracolumbar axis. We found significant differences between thoracic and lumbar cortical representations under vibrotactile stimulation at 80 Hz, but not at 20 Hz (Fig. 3). Specifically, at 80 Hz we found greater activation to upper, relative to lower back stimulation, in the right inferior parietal cortex extending into the superior temporal sulcus, as well as in bilateral regions of primary somatosensory cortex (Table 3). Atlas comparisons revealed the peak voxel activations of these latter clusters to be located in regions of high probabilistic overlap between multiple S1 sub-regions, with the left hemisphere result slightly favouring BA3b (40%, with 38% probability for BA3a and 29% for BA2) and the right hemisphere somewhat favouring BA2 (38%, with 33% for BA3a and 29% for BA3b). For the inverse contrast (80 Hz: lumbar > thoracic), however, a single significant cluster with its peak voxel located with strong probability (76%) in the right superior parietal lobule was detected (Table 3; an equivalent effect in the left hemisphere did not reach the cluster thresholding level required for statistical significance). No significant differences were identified for either of these contrasts in the 20 Hz stimulation condition. The results from our exploratory analysis of individual stimulation locations revealed some evidence of a representational progression along the thoracolumbar axis of primary somatosensory cortex (Fig. 4; Supplementary Figs. S1 and S2), where peak statistical activations and local maxima (see Tables in Supplementary Materials) tended to be represented in more ventral and anterior regions under upper back (thoracic) stimulation and in more dorsal and posterior regions under lower back (lumbar) stimulation.

**Figure 3.**
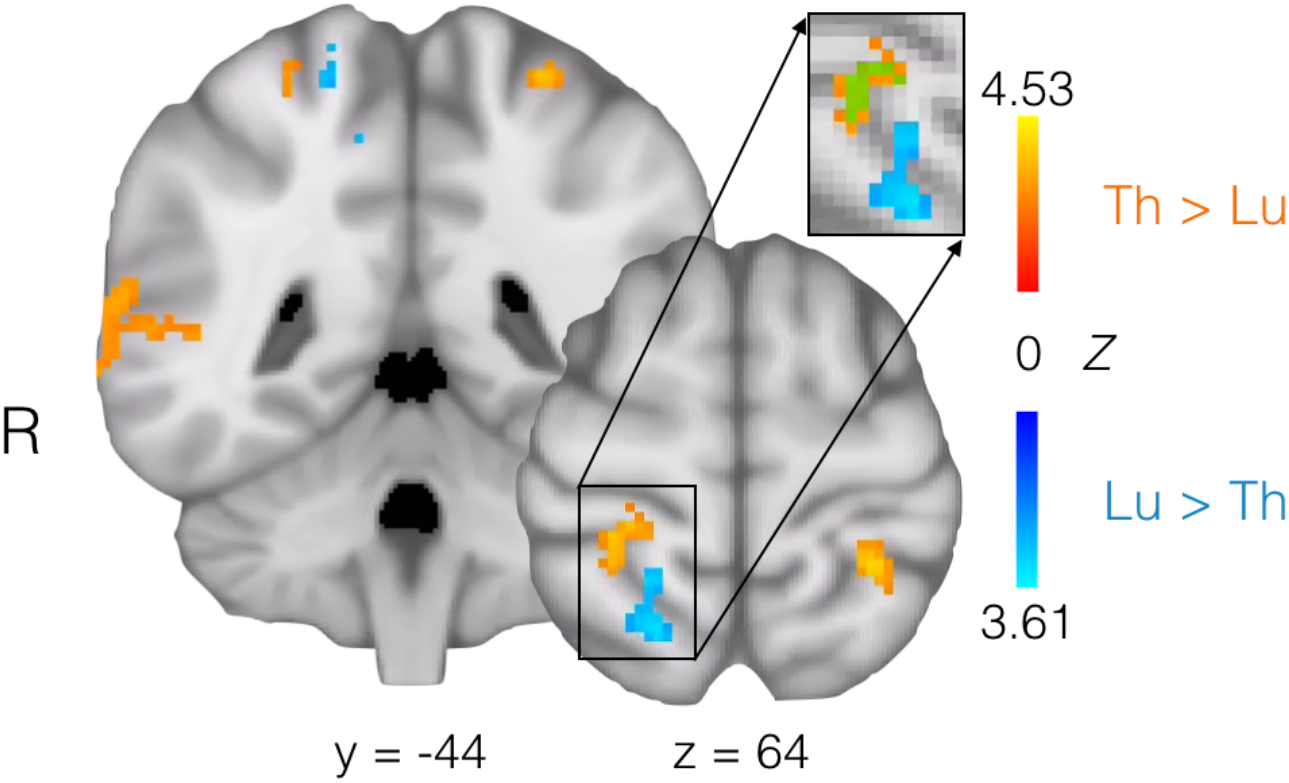
Brain regions displaying differential somatotopy, in terms of functional activation to proprioceptive (80 Hz) vibrotactile stimulation, depending on the location on the back. Red-yellow *Z*-statistic map and colour bar denote significantly increased activation to thoracic (upper back), relative to lumbar (lower back), stimulation (cluster *Z* > 2.3; p < 0.05; FWE-corrected; inset: green overlay with reduced opacity denotes significant equivalent cluster at a more conservative threshold of *Z* > 2.81); blue-cyan denotes a significant inverse (lumbar > thoracic) effect. Statistical maps are overlaid on a background image of a T1 template in MNI152 standard space. R = right hemisphere. NB: Nearby regions of significant activation for a given contrast can appear spatially distinct on the representative slices chosen for visualisation but still form part of the same spatially contiguous cluster.

**Table 3.** Brain regions significantly more activated under thoracic, relative to lumbar, 80 Hz proprioceptive vibrotactile stimulation of the back and vice versa. Three regional statistical maps displayed significantly greater activation (cluster *Z* > 2.3; p < 0.05; FWE-corrected; values in parentheses denote significant equivalent cluster information at a more conservative threshold of *Z* > 2.81) during thoracic stimulation than during lumbar simulation, while one showed the opposite effect. Statistical and anatomical co-ordinate information (relative to the MNI152 standard space template) is provided. Th = thoracic; Lu = lumbar; R = right hemisphere; L = left hemisphere; S1 = primary somatosensory cortex; SPL = superior parietal lobule; STS = superior temporal sulcus; BA = Brodmann area.

| Contrast | Voxels | P | Z-max | x (mm) | y (mm) | z (mm) | Anatomical loci |
| --- | --- | --- | --- | --- | --- | --- | --- |
| Th > Lu | 200 | < 0.001 | 3.26 | 64 | -46 | 16 | R angular & supramarginal gyri/STS |
|  | 153 | 0.003 | 4.53 | -22 | -36 | 54 | L S1 BA3b & BA3b/BA2 |
|  | 148<br>(62) | 0.004<br>(0.009) | 3.82 | 24 | -34 | 50 | R S1 BA2 & BA3a/BA3b |
| Lu > Th | 165 | 0.002 | 3.61 | 18 | -48 | 70 | R SPL |

**Figure 4.**
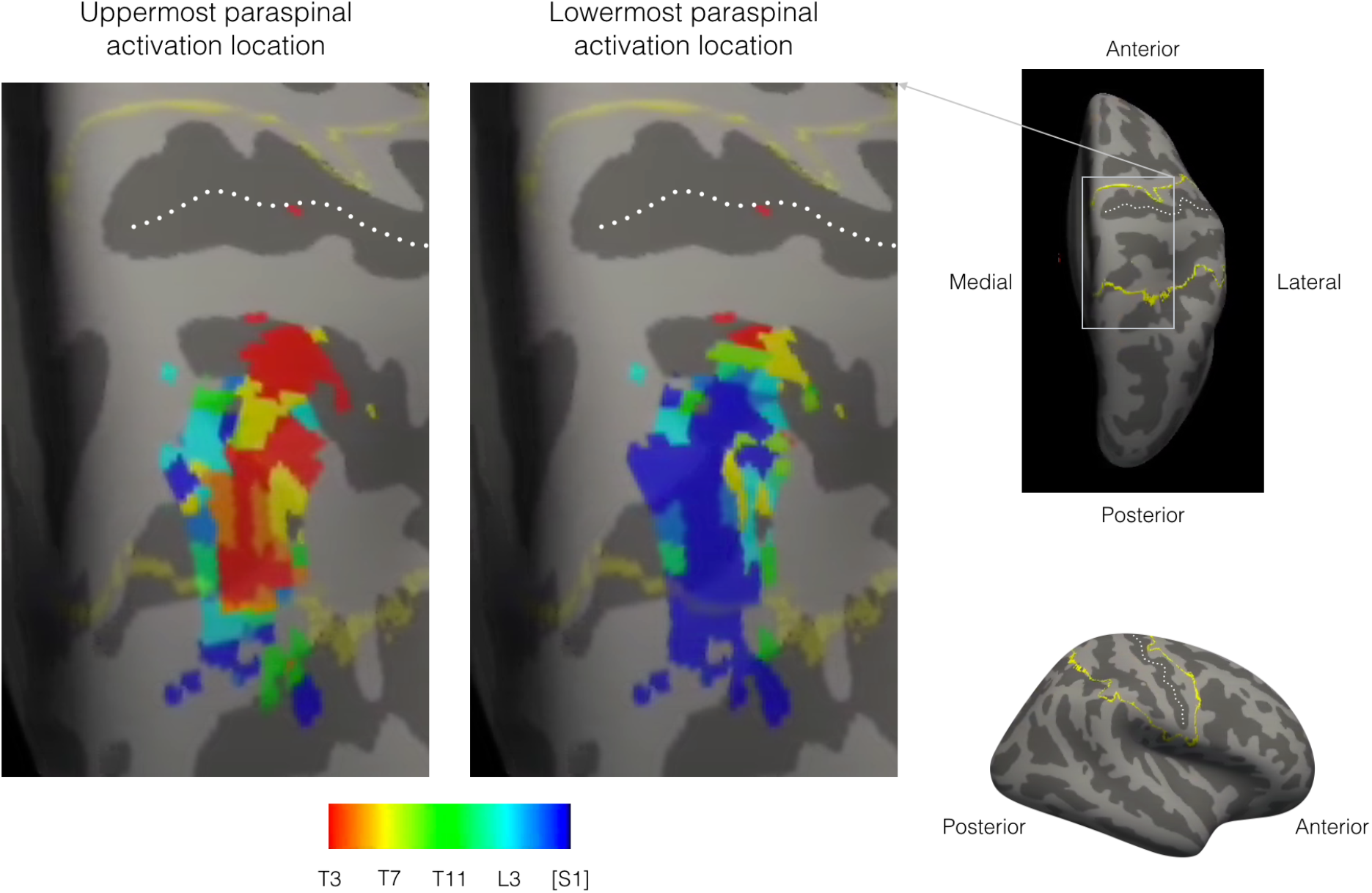
Discrete but overlapping somatotopy of upper and lower back is visible along the thoracolumbar axis. Suprathreshold activations (cluster *Z* > 2.3; p < 0.05; FWE-corrected) were binarized and indexed numerically from 1-9 according to vibration unit paraspinal location (see Fig. 1 and also Supplementary Fig. S2), concatenated across unit locations (using *fslmerge*), collated according to which unit was either the uppermost or the lowermost that evoked significant activation at a given voxel (using *fslmaths*) and then transformed (using the FreeSurfer tool *mri_vol2surf*) to an ‘inflated’ right hemisphere cortical surface template in MNI standard space. Colour bar represents which paraspinal location is either the uppermost or lowermost to significantly activate a given location in inflated surface space, ranging from T3 (red), to sacral[S1] (dark blue). A mask of primary somatosensory cortex (S1) comprising BA1, BA2 and BA3a/3b – based on unthresholded and then binarized and border-approximated (using *fslmaths*) Juelich Histological Atlas probabilistic maps – was also transformed to surface space and is displayed, for neuroanatomical orientation purposes, as a rough yellow boundary line: (i) overlaid with reduced opacity on the above-described uppermost/lowermost unit projection maps (or ‘paraspinal extremity-of-activation indices’), which are presented at a ‘zoomed in’ aerial posterior viewpoint, with the hemispheric surface angled 60º upwards from the anterior; (ii) overlaid at full opacity only on the surface template at the same viewpoint, but from a wide view (not zoomed in); and (iii) from a lateral wide viewpoint without any elevation. Dark grey regions of the surface template denote cortical sulci, while light grey denotes gyri. The central sulcus has been demarcated approximately as a dotted white line. The upper back predominantly appears to be represented more ventrally and anteriorly and the lower back predominantly represented more dorsally and posteriorly in S1 (with the middle back representation predominantly positioned in between the two).

## 4. Discussion

We present a first neurobiological validation of the novel ‘pneuVID’ method for mapping cortical representations of paraspinal afferent inputs. In particular we sought to delineate the neural representations of high- (80 Hz) versus low- (20 Hz) frequency paraspinal stimulation and the respective contributions of different mechanoreceptor types. Additionally, we attempted to provide an initial indication of the cortical somatotopy of multiple paraspinal locations across the thoracolumbar axis. The results suggest a spatial delineation of the functional representations of both paraspinal high-/low-frequency stimulation and upper/lower back sensory input across the span of the primary somatosensory and, to some extent, primary motor cortex. The pneuVID methodology may therefore serve as a promising tool for future investigations of detailed cortical maps of paraspinal afferent input and their potential alterations in neurological disorders or clinical conditions such as chronic LBP.

Firstly, we found distinct patterns of cortical activation representing high-frequency and low-frequency vibratory stimulations using pneuVID. In S1, peak representations of high-frequency stimulation were localised with a higher probability in region BA3a (on average 47%; BA3b = 10%), while activation of low-frequency stimulation was pinpointed to a greater extent in BA3b (average 28%; BA3a < 1%). This observation supports our hypothesis of spatial differences between cortical representations of low- and high-frequency vibration-evoked sensory information. Furthermore, the observed pattern regarding BA3a/3b activity could be explained by differential mechanotransduction generated by the two vibration frequencies. Neurons in BA3b respond primarily to cutaneous stimulation whereas neurons in BA3a exhibit mainly proprioceptive responses that are generated by joint, muscle and some skin mechanoreceptors, with the muscle spindle receptors providing the most important input for position and movement sense (i.e., kinaesthesia; for review see, e.g., Delhaye et al., 2018, although see also Kim et al., 2015). Higher-frequency, 80 Hz vibration is within the optimal bandwidth for stimulating primary (Ia) muscle spindle afferents (Roll and Vedel, 1982; Vedel and Roll, 1982; Brumagne et al., 2000; Seizova-Cajic et al., 2007) but may also co-stimulate Pacinian corpuscles (bandwidth between 80-400 Hz), which have been shown to contribute to proprioceptive function through skin stretch (Weerakkody et al., 2007). Thus, the observed dominance of BA3a activity might be a result of proprioceptive information processing during 80 Hz stimulation of the respective mechanoreceptors (primarily muscle spindles). Conversely, the neural activation in BA3b might be dominated by tactile information processing through stimulation of Meissner’s corpuscles during 20 Hz vibration. This is in line with prior findings detailed in the primate literature regarding the modality-specificity of equivalent hand and limb somatosensory perception (Garraghty et al., 1990; Romo et al., 1998; Naito et al., 1999; Delhaye et al., 2018). It must be noted, however, that experimental testing of the proprioceptive system in humans is often performed through the assessment of movement illusions (which were not systematically assessed in the current study). However, movement illusions are not easy to elicit in a consistent manner and are heavily dependent on the experimental context and the visual and tactile information processing during vibration (Taylor et al., 2017). Furthermore, vibration has been shown to elicit similar neural patterns to those associated with kinaesthesia in sensorimotor cortices, during both the perception and the absence of illusory movements (Schneider et al., 2021). As such, the perception of movement illusions cannot easily be considered as a mandatory prerequisite for the activation of proprioceptive systems.

In addition, peak activations were probabilistically localised more in primary motor cortex region BA4p under high-frequency stimulation (average 16.5%; BA4a < 1%) and in BA4a (40.3%; BA4p = 8.3%) under low-frequency stimulation. These findings of somatosensory information processing in motor cortical areas are in line with prior human and animal evidence in the literature, as it is known both that M1 activation is necessary for the somatosensory experience associated with movement of body parts such as the hand (Naito et al., 2002) and that cutaneous and proprioceptive inputs have direct pathways to M1 mediated through the spinothalamic and dorsal column tract, respectively (Asanuma et al., 1980; Avanzino et al., 2014). Furthermore, the observations in M1 appear to follow on from those in S1 as region BA4p, also known as “New M1”, is understood to be a phylogenetically newer evolutionary manifestation than region BA4a (“Old M1”), with the former being identified as specifically attuned to the complex processing of skilled motor behaviours characteristic of higher-order primates due to the prevalence of direct, monosynaptic connections between cortico-motor cells and muscle-adjacent motor neurons (Rathelot and Strick, 2009). This can be interpreted as mirroring the observed dissociations in S1 as area BA3a, which displayed a vibrotactile stimulation-related activation pattern more similar to BA4p than BA4a, is also understood to be a phylogenetically newer region than BA3b (Kaas, 2004). It is reasonable to assume that such cross-modal evolutionary development, integrating proprioceptive processing within newer sub-regions of somatosensory and motor cortex, has contributed to the amplification of complex abilities in humans and other higher order primates. In the current work we extend this framework of cross-modal integration to suggest that it applies to the human back and not only to bodily regions typically associated with more complex motor control and somatosensory perception, such as the hand. Further support for an increased complexity of the representations of spinal sensorimotor control in higher-order cortical regions has previously been provided by transcranial magnetic stimulation evidence of distinct motor cortical locations being associated with the excitation of deeper, relative to more superficial, paraspinal muscle fibres at the same lumbar location, again underlining the human ability for highly finely-graded control of these muscles (Tsao et al., 2011a). Interestingly, this complexity is thought to be central to our ability to adapt our movements and perceptions to the experience of pain (Hodges and Tucker, 2011; Tsao et al., 2011b; Elgueta-Cancino et al., 2021).

Secondly, we have expanded upon the work pioneered by Penfield (Penfield, 1947) and continued by others (e.g., Boendermaker et al., 2014; Roux et al., 2018) in mapping the functional topography of human somatosensory cortex, by providing additional information about fine-grained somatotopy of the back. Previously, very little has been established regarding the functional spatial organization of S1 with respect to the proprioceptive and other somatosensory representations of the back. In brief, Penfield first identified representational locations of the hip and the shoulder on the convexity of the postcentral gyrus and drew the representation of the back between these two areas (Penfield, 1947), while, more recently, intra-cortical stimulation of BA1 in S1 has provided confirmatory evidence that thoracic and abdominal neuroanatomical representations are, indeed, located between those of the hip and the shoulder (Roux et al., 2018), although the cortical somatotopic representation of the back along the thoracolumbar axis has remained unclear. Interestingly, a shifted representation of tactile stimuli applied to the back, away from the typically more lateral S1 regions and towards the midline, has been observed using magnetencephalography (MEG) in a small group of chronic LBP patients (Flor et al., 1997). In the current study, using fMRI, we were able to image S1 activation and other relevant brain regions at a higher spatial resolution, including information about cortical targets of different sensory afferents from different paraspinal locations. At a coarse, but statistically highly-powered level, we found high-frequency stimulation of the upper back, including thoracic areas, to activate regions of the inferior parietal cortex (angular and supramarginal gyri) and the anterior primary somatosensory cortex (BA2, BA3a and BA3b), while lower back stimulation including lumbar areas activated more superior and posterior regions of parietal cortex (superior parietal lobule area “5L”). At a finer-grained – but ultimately less statistically well-powered – level, when considering neuronal activations to stimulation of individual bilateral paraspinal locations, we have further confirmed this directional somatotopy during high-frequency stimulation (see Fig. 4 and Supplementary Materials). Regarding the apparent right hemisphere lateralisation of these above somatotopic observations, the present data provide limited scope with which to speculate about a hemispheric specialisation of function, given the fairly modest sample size. However, a precedent for this finding has perhaps previously been set in terms of highlighting the dominance of the right hemisphere in the perception of limb movement (Naito et al., 2005), a phenomenon that we contend may generalise to other areas of the somatosensory homunculus, including the back. In addition, it is worth noting that we were only able to explore and discuss differential somatotopic representations for high-frequency stimulation representations in detail, as for the majority of single-location low-frequency stimulation conditions no significant clusters were identified (in the 80 Hz condition stimulation of each paraspinal location was associated with at least one significant cluster). This is likely to be a corollary of the small sample size, which should be expanded upon in future neuroimaging studies attempting to investigate fine-grained tactile-perceptual representations of the back.

Further to the limitations already mentioned, it is important to discuss the possibility that our fMRI findings might instead specifically represent processes of differentiation between and encoding of (high and low) vibrotactile stimulation frequencies. Unfortunately, as we stimulated nine paraspinal locations at just two separate frequencies, in order to elevate statistical power for each condition without extending scan time to an uncomfortable length for participants, we are unable to test the frequency-dependence of representations in a more directly parametric manner using the current data. However, prior evidence has revealed that frequency-dependent representations of vibrotactile stimulation of the human index finger are more densely located in the lateral sulcus of S2 than in S1 or other regions of S2 (Chung et al., 2013), or in highly lateral regions of somatosensory cortex (reported as S1 but with peak coordinates located most probabilistically in S2 and the inferior parietal lobule) extending into the supramarginal gyrus (Kim et al., 2016). Therefore, it is possible that our findings, which centred around (comparatively medial regions of) S1, are more indicative of mechanoreceptor differentiation than of simple frequency encoding differences. Either way, it is likely that the differences between cellular processes distinguishing tactile/proprioceptive and frequency-encoding representations in S1 are too microstructural and interwoven in nature to be detected using whole-brain fMRI. Instead, it may only be currently feasible to delineate these processes using direct electrical recording techniques of small populations of neurons, above the limits of spatial resolution presently achievable with fMRI. However, progress in this area with fMRI may be expedited using computationally-inspired models of mechanistic spatial or frequency representation, for example with multi-voxel pattern analysis such as representational similarity analysis (Ejaz et al., 2015; Kim et al., 2016). In addition, higher field (e.g., 7 Tesla) MRI acquisitions may enable increased spatial delineation of different mechanistic processes due to the higher spatial resolutions they are able to achieve with fMRI, including at the level of different cortical layers: so-called “laminar fMRI” (Lawrence et al., 2019; Stephan et al., 2019).

Finally, if 80 Hz stimulation represents proprioceptive information processing, as is speculated above, it is not fully clear which specific trunk muscle spindles are targeted by vibratory stimulation with pneuVID. The primary assumption is that superficial (longissimus and spinalis) muscles along the thoracolumbar axis are most affected in terms of their activation. However, 80 Hz pneuVID stimulation might also penetrate to deeper muscles such as the rotatores and multifidi muscles, which are particularly dense in muscle spindle fibres and therefore also involved in key aspects of trunk and back proprioception (Boucher et al., 2015).

## 5. Conclusion

In sum, our findings highlight the potential for tools such as pneuVID to significantly extend previous work on mapping the cortical representation of paraspinal sensory input. Future combinations of the pneuVID technique with advanced computational and imaging methods should pave the way for new opportunities to map the cortical landscape of the human back in high detail and to elucidate its role in the development and maintenance of chronic LBP. Ultimately, this might provide promising neuroimaging-based outcomes with which to test the potential therapeutic effect of individualized therapies such as motor control exercises and how they compare to other treatment approaches (e.g., neuromodulation).

## 7. Data Availability Statement

The data that support the findings of this study are available from the corresponding author upon reasonable request. The data are not publicly available due to privacy or ethical restrictions.

## Acknowledgments

We thank Magdalena Suter for help with data collection and recruitment of participants. This research was supported by the Swiss National Science Foundation (SNSF, Bern, Switzerland).

