## Supplementary Materials for "In the back of your mind: Cortical mapping of paraspinal afferent inputs"

### Supplementary Figure Legends

**Figure S1.** Discrete somatotopy of upper and lower back is visible along the thoracolumbar axis. In the centre diagonal, statistical maps representing activation to 80 Hz stimulation at the single uppermost (orange-yellow), single precise middle (green) and single lowest (blue) back stimulation points are overlaid, with reduced opacity to aid visualisation of overlap, on multiple sequential slices from a background image of a T1 template in MNI152 standard space, with a focus on the right (R) hemisphere. The activation map representing stimulation of the leg (mTS) is also presented (red) to emphasise the concordance with – and the protraction of – known somatotopy. In the columnar plots, columns represent the frequency with which statistical peaks and other local maxima of activated clusters are located at specific single-plane MNI co-ordinates in the y-plane (bottom left; *x*-axis denotes anterior-posterior cortical positioning) and the z-plane (top right; *x*-axis = ventral-dorsal cortical elevation) for the uppermost *four* (orange-yellow), the single middle (green) and the lowest *four* (blue) stimulation sites (NB: this configuration does not precisely match that used in the brain image overlays described above, which was ungrouped and pruned for clarity of visualisation); (*) denotes the planar co-ordinate location of each of the nine peak voxels. The upper back appears to be represented more ventrally and anteriorly and the lower back represented more dorsally and posteriorly in S1 (with the middle back representation positioned in between the two).

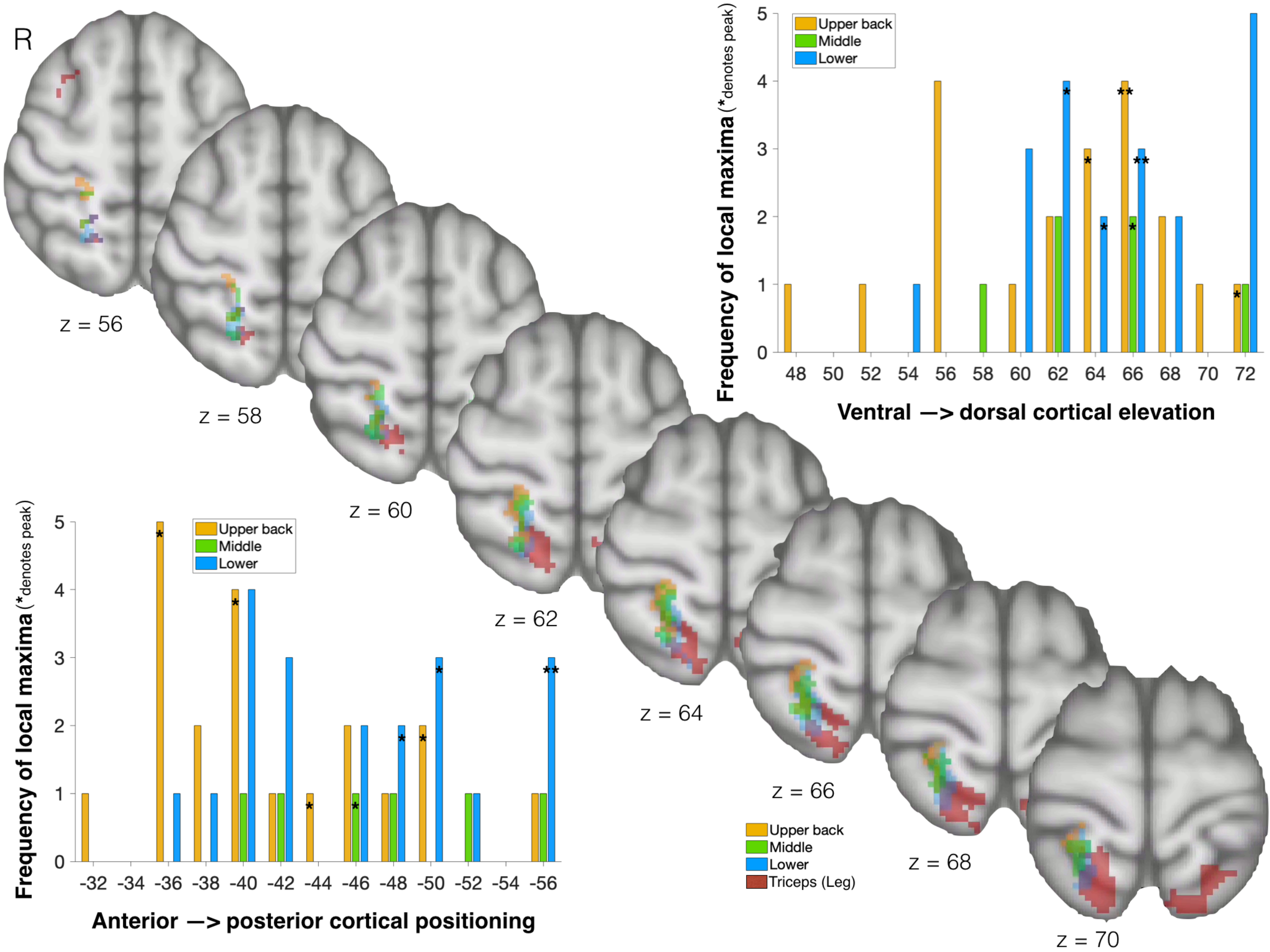

R
z = 56
z = 58
z = 60
z = 62
z = 64
z = 66
z = 68
z = 70
Upper back
Middle
Lower
Triceps (Leg)
Frequency of local maxima (*denotes peak)
Ventral —> dorsal cortical elevation
48 50 52 54 56 58 60 62 64 66 68 70 72
Anterior —> posterior cortical positioning
-32 -34 -36 -38 -40 -42 -44 -46 -48 -50 -52 -54 -56
0 1 2 3 4 5

**Figure S2.** Statistical maps of significant BOLD activity to high-frequency (80 Hz) bilateral stimulation of single pneuVID unit locations at (red) a less ($Z > 2.3$, $p < 0.05$; FWE-corrected) and (blue) a more ($Z > 2.81$, $p < 0.05$; FWE-corrected) conservative threshold. Figure panel numbers (1-9) denote, descending sequentially from upper to lower back regions, neural activity to vibrotactile stimulation at T3, T5, T7, T9, T11, L1, L3, L5 and sacral[S1]. Although not necessarily displaying a consistently obvious somatotopy, in general a displacement of activation moving from upper to lower back regions can be seen to map more posteriorly and dorsally in the primary somatosensory cortex. As not all units showed significant left hemisphere activation, here, for visual simplicity, only results projected on the right hemisphere (RH) are shown (cf. main text Fig. 4 to aid with spatial orientation). Suprathreshold activations were transformed to an 'inflated' RH cortical surface template in MNI standard space. Dark grey regions of the surface denote cortical sulci, while light grey denotes gyri.

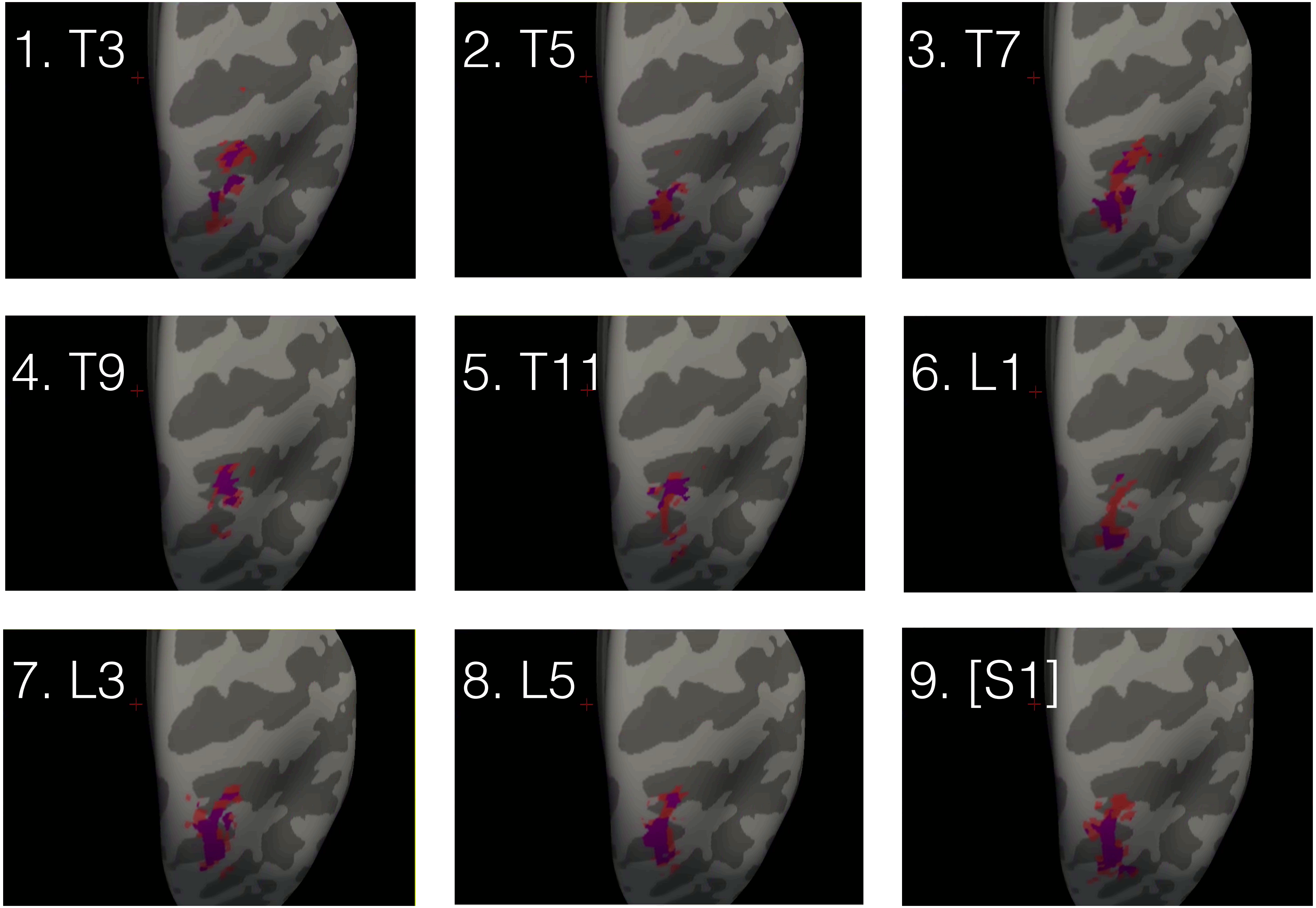
1. T3
2. T5
3. T7
4. T9
5. T11
6. L1
7. L3
8. L5
9. [S1]

T3

| Cluster Index | Z | x | y | z |
|---|---|---|---|---|
| 4 | 4.26 | -50 | -34 | 16 |
| 4 | 4.17 | -36 | -22 | 12 |
| 4 | 4.07 | -58 | -2 | 2 |
| 4 | 3.96 | -64 | -28 | 12 |
| 4 | 3.92 | -64 | -32 | 10 |
| 4 | 3.89 | -34 | -34 | 12 |
| 3 | 5.09 | 42 | -36 | 12 |
| 3 | 4.97 | 46 | -34 | 12 |
| 3 | 4.13 | 36 | -22 | 0 |
| 3 | 3.92 | 54 | -32 | 26 |
| 3 | 3.74 | 34 | -30 | 10 |
| 3 | 3.67 | 48 | -30 | 6 |
| 2 | 3.96 | 24 | -36 | 64 |
| 2 | 3.29 | 22 | -48 | 64 |
| 2 | 3.13 | 24 | -42 | 62 |
| 2 | 2.99 | 22 | -36 | 56 |
| 2 | 2.93 | 24 | -32 | 48 |
| 2 | 2.9 | 24 | -40 | 70 |
| 1 | 3.37 | 0 | -54 | 40 |
| 1 | 3.09 | -4 | -50 | 36 |
| 1 | 3.08 | 2 | -56 | 50 |
| 1 | 2.95 | -10 | -52 | 38 |
| 1 | 2.57 | 8 | -56 | 40 |
| 1 | 2.48 | 4 | -62 | 42 |

T5

| Cluster Index | Z | x | y | z |
|---|---|---|---|---|
| 5 | 4.39 | 44 | -36 | 14 |
| 5 | 3.62 | 52 | -34 | 14 |
| 5 | 3.39 | 52 | -36 | 18 |
| 5 | 3.21 | 58 | -28 | 16 |
| 5 | 3 | 54 | -30 | 24 |
| 4 | 4.27 | 38 | -20 | 12 |
| 4 | 4.07 | 42 | -16 | 4 |
| 4 | 3.93 | 38 | -22 | -4 |
| 4 | 3.86 | 36 | -24 | 0 |
| 4 | 2.97 | 42 | -10 | 2 |
| 4 | 2.91 | 30 | -22 | 0 |
| 3 | 3.34 | -56 | -30 | 10 |
| 3 | 3.14 | -64 | -26 | 12 |
| 3 | 3.11 | -42 | -34 | 12 |
| 3 | 3.1 | -42 | -42 | 20 |
| 3 | 3.02 | -58 | -22 | 12 |
| 3 | 3.02 | -64 | -20 | 12 |
| 2 | 4.7 | -36 | -20 | 12 |
| 2 | 3.68 | -50 | -16 | 8 |
| 2 | 3.24 | -34 | -24 | 2 |
| 2 | 2.57 | -38 | -18 | -4 |
| 2 | 2.37 | -56 | -16 | 12 |
| 1 | 3.7 | 24 | -50 | 66 |
| 1 | 3.44 | 24 | -36 | 66 |
| 1 | 3.1 | 22 | -56 | 56 |

T7

| Cluster Index | Z | x | y | z |
|---|---|---|---|---|
| 5 | 4.51 | -54 | -4 | 4 |
| 5 | 4.26 | -60 | -2 | 4 |
| 5 | 4.16 | -50 | -34 | 14 |
| 5 | 4.12 | -56 | -30 | 10 |
| 5 | 3.72 | -64 | -26 | 12 |
| 5 | 3.69 | -36 | -24 | -4 |
| 4 | 3.72 | 62 | -22 | 14 |
| 4 | 3.5 | 64 | -52 | 10 |
| 4 | 3.45 | 48 | -34 | 14 |
| 4 | 3.36 | 52 | -28 | 22 |
| 4 | 3.34 | 44 | -32 | 18 |
| 4 | 3.32 | 52 | -38 | 20 |
| 3 | 3.53 | 28 | -44 | 66 |
| 3 | 3.53 | 24 | -46 | 66 |
| 3 | 3.51 | 20 | -38 | 52 |
| 3 | 3.47 | 22 | -36 | 56 |
| 3 | 3.36 | 22 | -36 | 64 |
| 3 | 3.1 | 20 | -40 | 60 |
| 2 | 3.77 | -20 | -40 | 60 |
| 2 | 3.71 | -22 | -36 | 52 |
| 2 | 3.6 | -22 | -42 | 64 |
| 2 | 3.6 | -20 | -52 | 60 |
| 2 | 3.58 | -24 | -52 | 66 |
| 2 | 3.52 | -22 | -48 | 66 |
| 1 | 4.03 | 40 | -20 | 12 |
| 1 | 3.51 | 38 | -20 | -2 |
| 1 | 3.32 | 44 | -14 | 4 |
| 1 | 3.28 | 32 | -28 | 10 |
| 1 | 3.26 | 34 | -26 | 2 |

T9

| Cluster Index | Z | x | y | z |
|---|---|---|---|---|
| 4 | 4.1 | 44 | -36 | 12 |
| 4 | 4.08 | 40 | -36 | 14 |
| 4 | 3.72 | 34 | -24 | 2 |
| 4 | 3.72 | 36 | -26 | 12 |
| 4 | 3.63 | 36 | -22 | -2 |
| 4 | 3.58 | 38 | -18 | 4 |
| 3 | 4.28 | -36 | -22 | 12 |
| 3 | 3.75 | -34 | -32 | 12 |
| 3 | 3.64 | -34 | -24 | -2 |
| 3 | 3.63 | -30 | -28 | 6 |
| 3 | 3.6 | -48 | -36 | 14 |
| 3 | 3.32 | -66 | -22 | 10 |
| 2 | 3.96 | 22 | -40 | 72 |
| 2 | 3.85 | 22 | -40 | 62 |
| 2 | 3.28 | 20 | -38 | 56 |
| 2 | 3.28 | 24 | -50 | 68 |
| 2 | 3.25 | 24 | -46 | 68 |
| 1 | 4.06 | -58 | -2 | 4 |
| 1 | 3.96 | -56 | -8 | 4 |
| 1 | 2.52 | -58 | 6 | -2 |
| 1 | 2.46 | -64 | 4 | 8 |

T11

| Cluster Index | Z | x | y | z |
|---|---|---|---|---|
| 9 | 4.25 | 34 | -24 | 4 |
| 9 | 3.95 | 40 | -18 | 8 |
| 9 | 3.8 | 48 | -34 | 18 |
| 9 | 3.53 | 50 | -30 | 14 |
| 9 | 3.11 | 44 | -26 | 12 |
| 9 | 3.05 | 38 | -20 | -6 |
| 8 | 4.04 | -50 | -32 | 14 |
| 8 | 3.36 | -50 | -44 | 18 |
| 8 | 3.34 | -54 | -30 | 12 |
| 8 | 3.14 | -64 | -28 | 14 |
| 8 | 3.13 | -42 | -34 | 14 |
| 8 | 2.91 | -66 | -50 | 18 |
| 7 | 3.18 | -8 | -56 | 24 |
| 7 | 3.09 | -14 | -48 | 28 |
| 7 | 2.98 | -12 | -54 | 28 |
| 7 | 2.93 | -10 | -50 | 24 |
| 7 | 2.92 | -8 | -52 | 20 |
| 7 | 2.81 | -4 | -50 | 22 |
| 6 | 3.39 | -56 | -66 | 8 |
| 6 | 3.2 | -60 | -62 | 18 |
| 6 | 3.04 | -58 | -62 | 8 |
| 6 | 2.82 | -54 | -60 | 12 |
| 6 | 2.79 | -56 | -56 | 4 |
| 6 | 2.78 | -52 | -56 | 32 |
| 5 | 3.91 | 24 | -46 | 66 |
| 5 | 3.67 | 22 | -40 | 62 |
| 5 | 3.49 | 22 | -42 | 72 |
| 5 | 3.49 | 24 | -56 | 62 |
| 5 | 3.23 | 24 | -52 | 66 |
| 5 | 3.18 | 22 | -48 | 58 |
| 4 | 3.85 | -20 | -48 | 56 |
| 4 | 3.24 | -18 | -38 | 56 |
| 4 | 3.21 | -24 | -54 | 50 |
| 4 | 3.16 | -20 | -48 | 66 |
| 3 | 3.77 | -34 | -22 | 14 |
| 3 | 3.75 | -34 | -24 | 0 |
| 3 | 3.65 | -36 | -22 | -4 |
| 3 | 3.46 | -36 | -24 | 8 |
| 3 | 3.42 | -32 | -26 | 4 |
| 3 | 2.87 | -42 | -22 | 8 |
| 2 | 3.26 | -54 | 18 | 30 |
| 2 | 3.09 | -54 | 22 | 24 |
| 2 | 3.06 | -48 | 20 | 26 |
| 2 | 3.06 | -50 | 26 | 28 |
| 2 | 2.82 | -48 | 22 | 38 |
| 2 | 2.77 | -52 | 18 | 36 |
| 1 | 3.62 | -58 | 0 | 4 |
| 1 | 3.41 | -52 | -4 | 2 |
| 1 | 3.06 | -52 | -8 | 6 |
| 1 | 3.01 | -44 | -8 | 0 |

| | | | | |
|---|---|---|---|---|
| 1 | 2.96 | -56 | -8 | 4 |

L1

| Cluster Index | Z | x | y | z |
|---|---|---|---|---|
| 4 | 4.18 | 42 | -36 | 14 |
| 4 | 4.16 | 50 | -34 | 18 |
| 4 | 3.38 | 66 | -18 | 8 |
| 4 | 3.28 | 68 | -22 | 20 |
| 4 | 3.26 | 58 | -18 | 16 |
| 4 | 3.24 | 64 | -30 | 18 |
| 3 | 4.24 | -48 | -36 | 16 |
| 3 | 4.21 | -60 | -22 | 12 |
| 3 | 3.9 | -52 | -32 | 12 |
| 3 | 3.15 | -60 | -28 | 10 |
| 3 | 3.02 | -42 | -32 | 10 |
| 3 | 2.99 | -48 | -26 | 12 |
| 2 | 4.07 | 22 | -56 | 62 |
| 2 | 3.09 | 28 | -42 | 68 |
| 2 | 3.04 | 22 | -42 | 72 |
| 2 | 2.87 | 20 | -38 | 64 |
| 2 | 2.55 | 24 | -50 | 54 |
| 2 | 2.45 | 24 | -46 | 66 |
| 1 | 3.44 | 36 | -18 | 2 |
| 1 | 3.37 | 38 | -20 | 12 |
| 1 | 3.32 | 40 | -16 | 2 |
| 1 | 2.92 | 42 | -14 | 18 |
| 1 | 2.69 | 38 | -20 | -6 |
| 1 | 2.53 | 44 | -12 | 4 |

L3

| Cluster Index | Z | x | y | z |
|---|---|---|---|---|
| 5 | 4.48 | 22 | -48 | 66 |
| 5 | 3.61 | 20 | -52 | 60 |
| 5 | 3.53 | 22 | -40 | 60 |
| 5 | 3.36 | 22 | -40 | 72 |
| 4 | 4.13 | 36 | -22 | 4 |
| 4 | 4.08 | 38 | -20 | 12 |
| 4 | 3.7 | 46 | -14 | 16 |
| 4 | 3.1 | 54 | -20 | 20 |
| 4 | 2.85 | 56 | -18 | 24 |
| 4 | 2.5 | 60 | -20 | 16 |
| 3 | 3.48 | -48 | -64 | 6 |
| 3 | 3.4 | -42 | -60 | 6 |
| 3 | 3.31 | -58 | -64 | 8 |
| 3 | 2.9 | -52 | -58 | 10 |
| 2 | 3.93 | 42 | -38 | 12 |
| 2 | 3.78 | 44 | -30 | 18 |
| 2 | 2.71 | 46 | -26 | 18 |
| 2 | 2.63 | 54 | -38 | 24 |
| 2 | 2.51 | 50 | -36 | 26 |
| 2 | 2.44 | 52 | -28 | 22 |
| 1 | 4.1 | -54 | -2 | 4 |
| 1 | 3.15 | -60 | -4 | 4 |

L5

| Cluster Index | Z | x | y | z |
|---|---|---|---|---|
| 4 | 4 | 44 | -36 | 12 |
| 4 | 3.87 | 40 | -20 | 12 |
| 4 | 3.65 | 36 | -20 | 0 |
| 4 | 3.56 | 48 | -34 | 18 |
| 4 | 3.56 | 38 | -22 | -4 |
| 4 | 3.52 | 32 | -24 | 6 |
| 3 | 3.97 | -54 | -30 | 12 |
| 3 | 3.47 | -64 | -34 | 16 |
| 3 | 3.41 | -48 | -34 | 14 |
| 3 | 2.99 | -60 | -30 | 16 |
| 3 | 2.97 | -62 | -20 | 20 |
| 3 | 2.96 | -50 | -40 | 16 |
| 2 | 4.25 | 22 | -56 | 64 |
| 2 | 3.59 | 20 | -48 | 62 |
| 2 | 3.41 | 22 | -42 | 72 |
| 2 | 3.14 | 22 | -40 | 62 |
| 1 | 3.99 | -34 | -22 | 12 |
| 1 | 3.75 | -34 | -22 | 0 |
| 1 | 2.77 | -28 | -26 | 14 |
| 1 | 2.71 | -44 | -20 | 12 |
| 1 | 2.62 | -44 | -24 | 6 |
| 1 | 2.45 | -46 | -24 | 14 |

Sacral S1

| Cluster Index | Z | x | y | z |
|---|---|---|---|---|
| 5 | 4.66 | -48 | -36 | 14 |
| 5 | 4.6 | -34 | -22 | 10 |
| 5 | 4.21 | -64 | -34 | 14 |
| 5 | 4.19 | -32 | -24 | 4 |
| 5 | 4.18 | -66 | -22 | 12 |
| 5 | 3.99 | -36 | -18 | -2 |
| 4 | 4.39 | 38 | -20 | 8 |
| 4 | 4.23 | 38 | -20 | -4 |
| 4 | 4.14 | 64 | -26 | 18 |
| 4 | 4.02 | 38 | -20 | 12 |
| 4 | 4.02 | 46 | -34 | 18 |
| 4 | 4.01 | 38 | -20 | 2 |
| 3 | 3.39 | 0 | -72 | 26 |
| 3 | 3.33 | 2 | -58 | 12 |
| 3 | 3.33 | 0 | -66 | 18 |
| 3 | 3.25 | -2 | -52 | 6 |
| 3 | 3.23 | 2 | -54 | 24 |
| 3 | 3.19 | -2 | -60 | 24 |
| 2 | 4.14 | 20 | -50 | 66 |
| 2 | 4.08 | 22 | -56 | 62 |
| 2 | 3.32 | 22 | -40 | 72 |
| 2 | 3.27 | 20 | -50 | 72 |
| 2 | 3.24 | 22 | -36 | 68 |
| 2 | 3.12 | 20 | -46 | 60 |
| 1 | 3.78 | -60 | 0 | 2 |
| 1 | 3.09 | -56 | -4 | -2 |
| 1 | 3.07 | -48 | -8 | 4 |
| 1 | 2.99 | -48 | -4 | 0 |

mTS

| Cluster Index | Z | x | y | z |
|---|---|---|---|---|
| 6 | 5.02 | -36 | -22 | 6 |
| 6 | 3.74 | -56 | -20 | 12 |
| 6 | 3.52 | -44 | -6 | 4 |
| 6 | 3.36 | -56 | -28 | 10 |
| 6 | 3.29 | -64 | -32 | 20 |
| 6 | 3.28 | -48 | -26 | 12 |
| 5 | 4.49 | 38 | -20 | 8 |
| 5 | 4.44 | 32 | -28 | 10 |
| 5 | 4.06 | 34 | -24 | 2 |
| 5 | 3.87 | 38 | -22 | -4 |
| 5 | 3.85 | 46 | -36 | 14 |
| 5 | 3.47 | 58 | -18 | 8 |
| 4 | 4.56 | 18 | -60 | 62 |
| 4 | 3.87 | 14 | -54 | 70 |
| 4 | 3.19 | 12 | -54 | 64 |
| 4 | 2.98 | 16 | -42 | 72 |
| 4 | 2.86 | 22 | -48 | 64 |
| 4 | 2.85 | 10 | -64 | 66 |
| 3 | 4.37 | -16 | -62 | 68 |
| 3 | 3.72 | -10 | -58 | 66 |
| 3 | 3.47 | -6 | -56 | 64 |
| 3 | 2.87 | -20 | -48 | 68 |
| 3 | 2.49 | -28 | -50 | 68 |
| 2 | 3.49 | 38 | -76 | 34 |
| 2 | 3.11 | 42 | -68 | 40 |
| 2 | 3.09 | 46 | -66 | 38 |
| 2 | 2.95 | 50 | -60 | 32 |
| 2 | 2.92 | 40 | -64 | 38 |
| 2 | 2.81 | 48 | -72 | 34 |
| 1 | 3.41 | 26 | 16 | 42 |
| 1 | 3.25 | 32 | 10 | 52 |
| 1 | 3.04 | 34 | 14 | 50 |
| 1 | 2.75 | 40 | 12 | 50 |
